# HEPLISAV-B Breaks Immune Tolerance and Induces HBV Control via CD4 T Cell-Dependent Mechanisms in a Chronic Hepatitis B Mouse Model

**DOI:** 10.64898/2026.03.13.711721

**Authors:** James Ahodantin, Jiapeng Wu, Masaya Funaki, Lydia Tang, Shyamasundaran Kottilil, Lishan Su

## Abstract

**Background:** Chronic hepatitis B virus (HBV) infection (CHB) affects nearly 300 million individuals globally and remains incurable with current antiviral therapies, which suppress viral replication but rarely achieve functional cure defined by sustained loss of hepatitis B surface antigen (HBsAg). CHB is characterized by profound virus-induced immune tolerance that limits the efficacy of conventional therapeutic vaccination strategies.

**Objective:** To evaluate the therapeutic efficacy and immunological mechanisms of HEPLISAV-B, a CpG-1018–adjuvanted HBsAg vaccine, in breaking immune tolerance and inducing functional cure–like responses in a murine model of CHB.

**Design:** Using the adeno-associated virus-HBV (AAV-HBV) mouse model, mice with high levels of persistent HBV viremia were vaccinated with two doses of HEPLISAV-B. Virological outcomes in the blood and liver, immune responses and mechanisms were assessed.

**Results:** HEPLISAV-B induced rapid and durable HBsAg clearance, markedly reduced circulating and intrahepatic HBV DNA and RNA, and suppressed viral replication without hepatocellular injury. Vaccination elicited robust, sustained anti-HBs IgG1 and IgA responses, enhanced HBsAg-specific T and B cell immunity, reduced CD4 regulatory T cells, and decreased PD-1 expression on CD4 T cells. Therapeutic efficacy was strictly dependent on CD4 T cells and the CD40/CD40L signaling pathway, but independent of CD8 T cells, indicating a CD4-driven, non-cytolytic antiviral mechanism critical for HEPLISAV-B induced HBV control.

**Conclusion:** HEPLISAV-B effectively breaks HBV-induced immune tolerance and restores coordinated antiviral immunity through a CD4 T cell–/CD40L-dependent pathway. The findings support its potential as a therapeutic vaccine in CHB patients.

**Key messages:** *What is already known on this topic:* Chronic HBV infection is marked by profound virus-induced immune tolerance, current antiviral therapies and vaccines fail to reliably induce HBsAg loss or restore effective antiviral immunity, highlighting the need for immune-based therapeutic strategies.

*What this study adds:* This study demonstrates that the clinically approved vaccine HEPLISAV-B can break HBV immune tolerance in a chronic HBV mouse model, inducing durable HBsAg clearance and anti-HBs immunity, non-cytolytic depletion of intrahepatic HBV DNA, through a mechanism strictly dependent on CD4 T cells and CD40/CD40L signaling.

*How this study might affect research, practice or policy:* These findings defined a CD4 T cell-CD40L/CD40 axis that is critical in CHB functional cure, and support testing HEPLISAV-B as a therapeutic vaccine in CHB patients.

Graphical Abstract

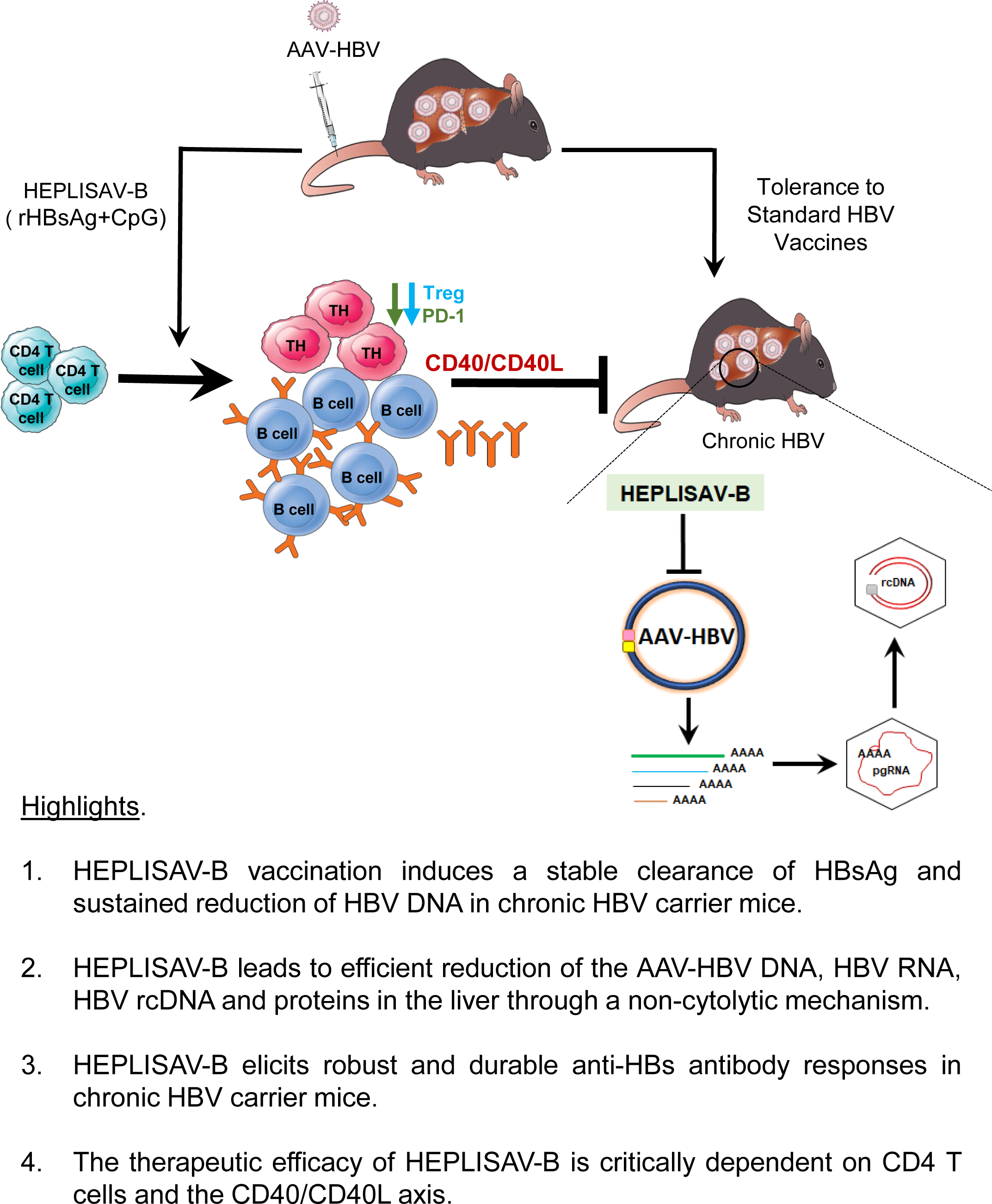

## Introduction

Chronic hepatitis B virus (HBV) infection remains a major global health issue, affecting approximately 296 million people worldwide^1^. It can lead to severe liver complications, including cirrhosis and hepatocellular carcinoma. Current nucleos(t)ide analogue treatments, while effective in suppressing viral replication, rarely achieve a functional cure, defined as sustained hepatitis B surface antigen (HBsAg) and HBV DNA loss after discontinuation of HBV medications^2, 3^.

People with resolved acute HBV infection typically exhibit robust systemic antibody, CD4, and CD8 T-cell responses against HBV^4–6^. In contrast, individuals with chronic HBV infection fail to produce effective anti-HBs antibodies^7^ and have dysfunctional HBV-specific CD4 and CD8 T cells, resulting in HBV-specific immune tolerance^8^, characterized by high HBV viral load with minimal liver inflammation. High levels of systemic viral antigens induce immune tolerance, impairing dendritic cell, natural killer cells, and T- and B-cell functions, thereby contributing to HBV persistence^9–14^. Overcoming this tolerance to elicit effective anti-HBV immune responses remains a major challenge in developing HBV therapeutic vaccines. Therefore, novel treatments, including therapeutic vaccines capable of boosting anti-HBV immunity, are urgently needed to effectively eradicate chronic HBV infection and improve patients’ lives.

Our group and others have shown that optimizing immune adjuvants is a promising strategy to overcome HBV-induced immune tolerance^15–24^. CpG oligodeoxynucleotides (CpG ODNs), potent TLR9 agonists, have shown powerful adjuvant properties that kickstart a strong immune response capable of addressing the inherent immune hypo-responsiveness in chronic HBV patients (CHB). By activating and maturing Antigen Presenting Cells (especially pDCs), TLR9 agonists ensure that co-administered HBV antigens are efficiently processed and presented to naive and exhausted T cells, thereby initiating or boosting a specific anti-HBV immune response^25, 26^.

We have previously demonstrated that CpG oligodeoxynucleotide 1826 (CpG-1826) as an adjuvant to HBV vaccines shows promise in breaking immune tolerance and achieving viral clearance in the AAV-HBV mouse model^15^. However, that study focused on short-term effects, leaving the long-term durability of the immune response, the risk of viral rebound, and potential adverse effects largely unexplored. Furthermore, CpG-1826 vaccination failed to elicit responses in mice with high HBV viremia^15^.

CpG-1018, a Dynavax-developed CpG, is used as an adjuvant in HEPLISAV-B, the novel FDA-approved vaccine for inducing protective HBV immunity in population who fail to respond to conventional HBV vaccines such as Engerix-B^27, 28^. HEPLISAV-B vaccine has demonstrated superior immunogenicity compared to alum-adjuvanted vaccines, even in populations with traditionally poor vaccine responses (e.g., people with HIV, diabetes, or chronic kidney disease)^29–31^. In clinical trials, CpG 1018-adjuvanted vaccines induced higher seroconversion rates, faster immune responses, and stronger T and B cell memory^29–31^.

In the present study, we evaluated the therapeutic efficacy of the HEPLISAV-B vaccine in a murine model of chronic HBV infection. In contrast to conventional HBsAg vaccination, which failed to overcome established immune tolerance and exhibited no antiviral activity, HEPLISAV-B induced rapid and sustained clearance of circulating HBsAg, markedly reduced intrahepatic HBV DNA, and suppressed viral replication through a predominantly non-cytolytic mechanism.

HEPLISAV-B also elicited robust and durable anti-HBs antibody responses and was associated with a reduction of CD4 regulatory T cells and decreased PD-1 expression on CD4 T cells. The therapeutic effect of HEPLISAV-B was dependent on CD4 T cells and CD40L, but not CD8 T cells. Together, these findings establish HEPLISAV-B as a potent immunotherapeutic capable of breaking CHB immune tolerance and inducing coordinated antiviral immunity, supporting its potential use as a therapeutic vaccine for achieving a functional cure in patients with chronic HBV infection.

## Materials and Methods

### Animals’ husbandry and Treatments

Male C57BL/6 mice were housed under standard conditions (23 ± 2 °C, 12-h light/dark cycle, ad libitum food and water). All procedures were approved by the University of Maryland School of Medicine IACUC (AUP-00000478). Eight- to nine-week-old mice were intravenously injected with recombinant AAV-HBV (5×10^10^ genome copies/mouse), and plasma HBV viremia and HBsAg levels were monitored every 2-4 weeks for up to 24 weeks high HBV viremia^15^.

In the Montanide/mIL12 immunization study, HBV-positive and uninfected control mice received two subcutaneous immunizations (4 weeks apart) of HBsAg (20 µg) emulsified in Montanide ISA 720 VG (25% v/v), together with intramuscular injections of pVAX1-coHBcAg (100 µg) and pORF-mIL-12 (20 µg) DNA per mouse. In the HEPLISAV-B titration experiment, mock-infected and HBV-positive mice received two intramuscular doses of HEPLISAV-B (4 weeks apart) at varying rHBsAg/CpG doses (0.08/12, 0.4/60, or 2/300 µg). To assess immune mechanisms, HEPLISAV-B–treated HBV-positive mice were administered with anti-CD4 or anti-CD8 antibodies for T-cell depletion, or anti-CD40L antibody to block CD40L–CD40 signaling.

### HBV Viremia Analysis

HBV DNA was extracted from 25 µL of plasma using the QIAamp DNA Blood Mini Kit (Qiagen, #51106). Plasma HBV DNA levels were quantified by qPCR using Power SYBR Green PCR Master Mix (Applied Biosystems) on a CFX Opus 384 Real-Time PCR System (Bio-Rad) with HBV-specific mouse primers (Table 1). Absolute HBV DNA copy numbers were calculated using standard curves generated from an HBV genome–containing plasmid, as previously described^32^.

**Table 1.**
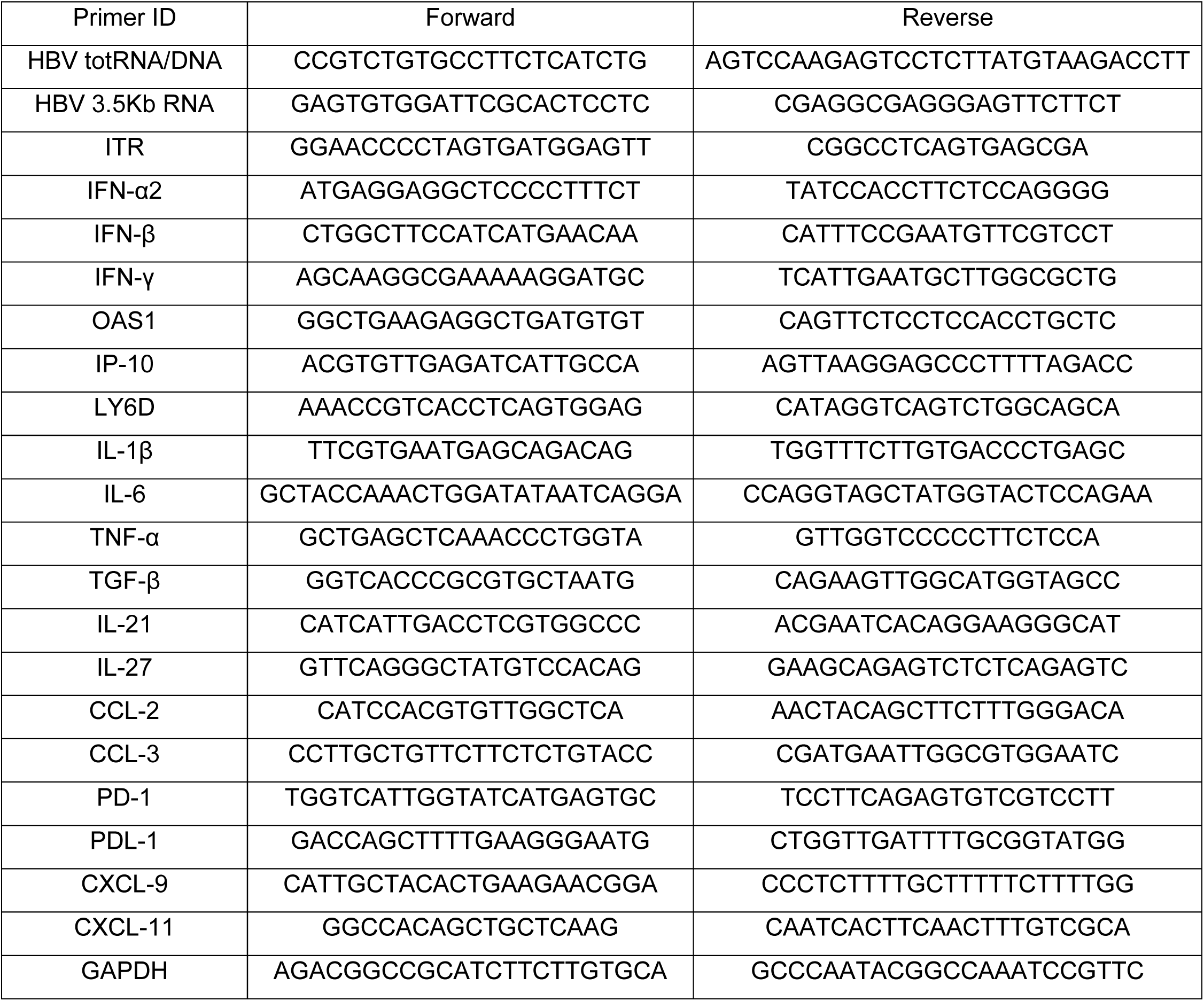
Primers sequences for qPCR/RT-qPCR.

### DNA/RNA Extraction and Real-Time Quantitative PCR

Total RNA was isolated from cells using the RNeasy Plus Kit and from liver tissue using QIAzol Lysis Reagent with gDNA Eliminator Solution (RNeasy Plus Universal Kit; QIAGEN) and quantified by NanoDrop. cDNA was synthesized from 0.2–2 µg of total RNA using random primers and SuperScript III (Invitrogen). Genomic DNA was extracted from 50 mg of liver tissue using the MasterPure™ Complete DNA and RNA Purification Kit (Biosearch Technologies). Diluted cDNA and DNA (1:10; 2 µL) were quantified by qPCR using Power SYBR Green PCR Master Mix (Applied Biosystems) on a CFX Opus 384 Real-Time PCR System (Bio-Rad) with mouse-specific primers (Table 1). Relative expression levels were calculated using the comparative Ct method, normalized to mouse GAPDH, and expressed relative to control animals^33–36^.

### Protein Extraction and Immunoblot

Total protein was extracted from snap-frozen liver tissue using RIPA buffer (Thermo Scientific) supplemented with protease and phosphatase inhibitor cocktails (Pierce) and quantified by BCA assay. Plasma (3 µL) and liver protein extracts (5–20 µg) were separated by SDS–PAGE (4–12%), transferred to nitrocellulose membranes, and probed with primary antibodies against HBs (1:1000; BIOSYNTH, #20-HR20) and HBc (1:1000; ZETA Corp, #Z2085RL-R). Horseradish peroxidase–conjugated anti-mouse or anti-rabbit secondary antibodies were used, and HRP-conjugated β-actin (1:5000; Sigma/Millipore, A3854-200 UL) served as a loading control. Signals were detected by ECL (Millipore or Invitrogen) using a Bio-Rad CCD camera and quantified with ImageJ software.

### Histology and Immunofluorescence Staining

For histological and immunofluorescence analyses, formalin-fixed, paraffin-embedded mouse liver sections (5 µm) were stained with hematoxylin and eosin (H&E) or subjected to antigen retrieval and immunostained with anti-HBs (BIOSYNTH, #20-HR20) and anti-HBc (ZETA Corp, #Z2085RL-R) antibodies, followed by Alexa Fluor 488–conjugated anti-rabbit secondary antibody (1:500; Invitrogen). Apoptotic cells were detected using a TUNEL assay kit (Roche) according to the manufacturer’s instructions. Nuclei were counterstained with Hoechst 33342 (Sigma), and images were acquired using a 20x objective.

### Immune Cells Isolation, Treatment and Flow Cytometry

Liver immune cells were isolated as previously described, with minor modifications.³⁶ Briefly, livers were perfused via the portal vein with wash buffer (PBS, 2% FBS, 1% penicillin/streptomycin, 0.1 mg/mL DNase I), minced using a gentleMACS Dissociator, and digested in Liver Digestion Medium (Thermo Fisher, #17703034) for 30 min at 37 °C. Cell suspensions were layered onto a 40%/80% Percoll gradient and centrifuged at 800 × g for 30 min at room temperature without brake. Immune cells were collected from the interphase, washed, and processed for downstream analyses. Splenocytes were prepared similarly, and red blood cells were lysed with ACK buffer (Invitrogen).

For ex vivo T-cell stimulation, cells were incubated with PMA and ionomycin for 8 h, with brefeldin A added during the final 6 h. Cells were blocked with anti-FcR (CD16/32; BioLegend) for 15 min, stained with fluorophore-conjugated antibodies for surface markers at 4 °C for 30 min, fixed and permeabilized using the FOXP3 Fixation/Permeabilization buffer (eBioscience), and subsequently stained for intracellular markers (Table 2). Samples were acquired on a spectral Aurora UV flow cytometer (Cytiva/Cytek).

**Table 2.**
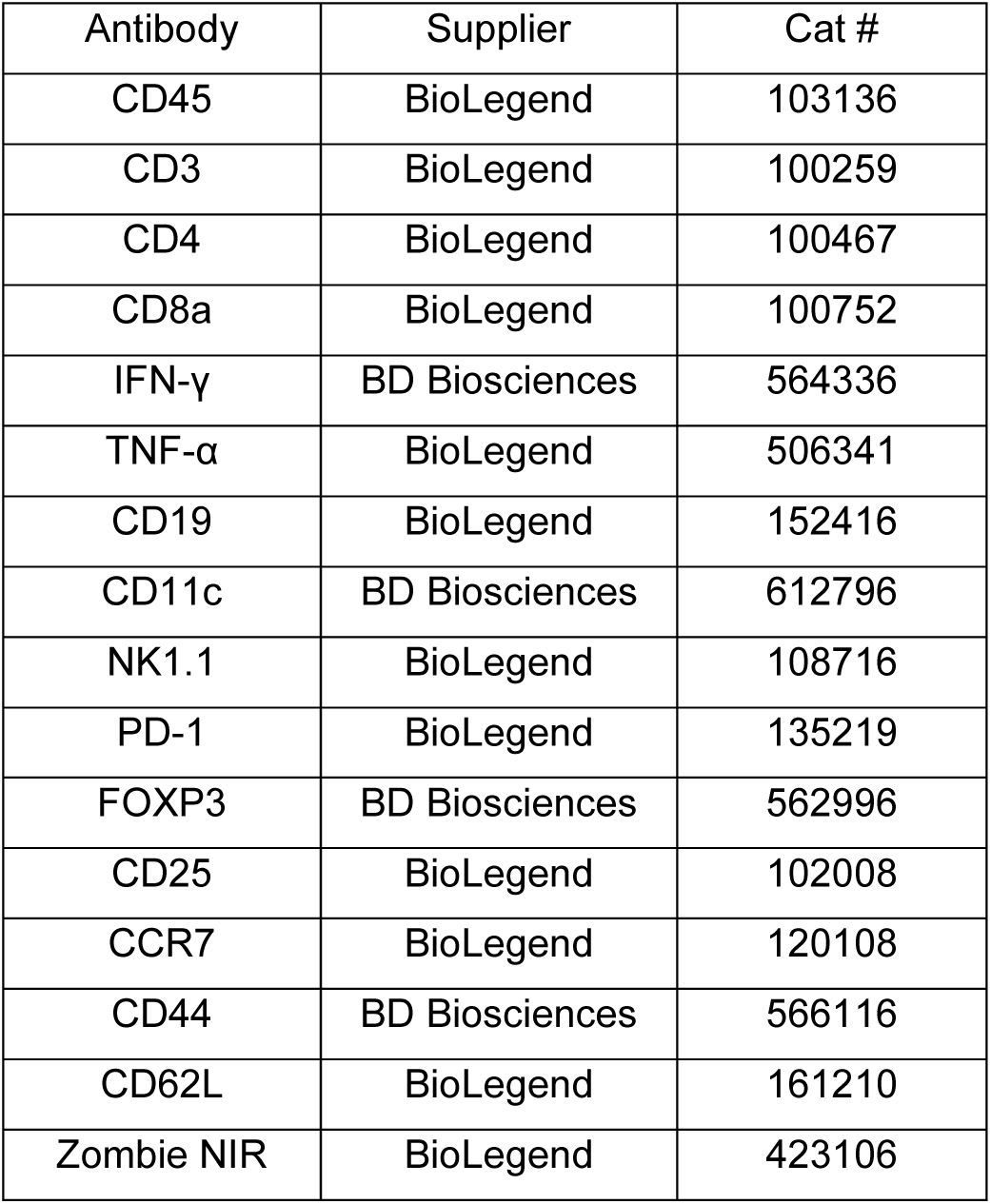
Flow Cytometry Antibodies.

### ELISPOT Assays

ELISPOT assays were performed on splenocytes isolated from mock- and AAV-HBV–infected mice, with or without HEPLISAV-B treatment. IFN-γ–producing cells were assessed after 48 h stimulation with HBsAg (20 µg/mL) in the presence of CD28/CD49d co-stimulatory antibodies (BD Biosciences, #347690); CD3/CD28 microbeads served as positive controls. For total and HBsAg-specific IgG detection, splenocytes were pre-stimulated with mouse B-Poly-S™ reagent for 5 days, then transferred to pre-activated ELISPOT plates coated with Igκ/λ or HBsAg (10 µg/mL) and incubated for 16 h. Assays were performed according to the manufacturer’s instructions, and plates were analyzed using a CTL ImmunoSpot® Analyzer and ImmunoSpot® Software, with quality control conducted by an experienced ELISPOT specialist.

### Quantification of Secreted Proteins by Enzyme Linked Immunosorbent Assay (ELISA)

Plasma levels of alanine aminotransferase (ALT; Elabscience®, #E-BC-K235-M), HBsAg (Alphadiagnostic International, #4110), HBeAg (Ig Biotechnology, SKUs CL18004 and CL0312-2), and mouse anti-HBs IgG (Alphadiagnostic International, #4210) were quantified by ELISA according to the manufacturers’ instructions. HBsAg-specific IgG1, IgG2b, and IgA were measured using HBsAg-coated plates in combination with mouse IgG1/2b and IgA ELISA kits (Thermo Fisher Scientific, #88-50410-22, #88-50430-22, and #88-50450-22).

### Statistical Analysis

Statistical significance was assessed by two-tailed unpaired Student t tests (used to analyze means in normally distributed populations) or one- and two-way analysis of variance (used to analyze ≥ 3 variables) using Prism 10 (GraphPad Software).

## Results

### HEPLISAV-B breaks tolerance and suppresses HBV production in chronic HBV-carrier mice

Consistent with prior reports that alum-adjuvanted Engerix-B fails to elicit anti-HBs in AAV-HBV carrier mice^27, 28^, Montanide ISA 720 VG with co-administration of HBcAg/IL-12 DNA induced no seroconversion nor reduced viremia (Figure S1A-G). HBsAg and HBcAg remained unchanged in liver (Figure S1E-G), confirming that conventional adjuvants do not overcome HBV-induced tolerance. We next evaluated HEPLISAV-B. Dose-escalation in HBV-naïve mice identified dose 3 (HBsAg/CpG-1018: 2 µg/300 µg) as optimal, inducing the highest sustained anti-HBs titers after two injections (Figure S2A-B). In chronic AAV-HBV mice vaccinated at 4 and 8 wpi (Figure 1A), HEPLISAV-B produced a 1,000-fold reduction in serum HBV DNA at 8-12 wpi (Figure 1B), with levels remaining significantly lower than controls through 24 wpi despite a partial rebound at 16 wpi. HBeAg declined significantly (Figure 1C), and, strikingly, HBsAg became undetectable, whereas it persisted in controls (Figure 1D). Immunoblotting confirmed marked depletion of HBsAg and HBcAg in plasma of vaccinated mice at 24 wpi (Figure 1E-G), while ALT remained unchanged (Figure 1H), indicating non-cytolytic viral control.

**Figure 1.**
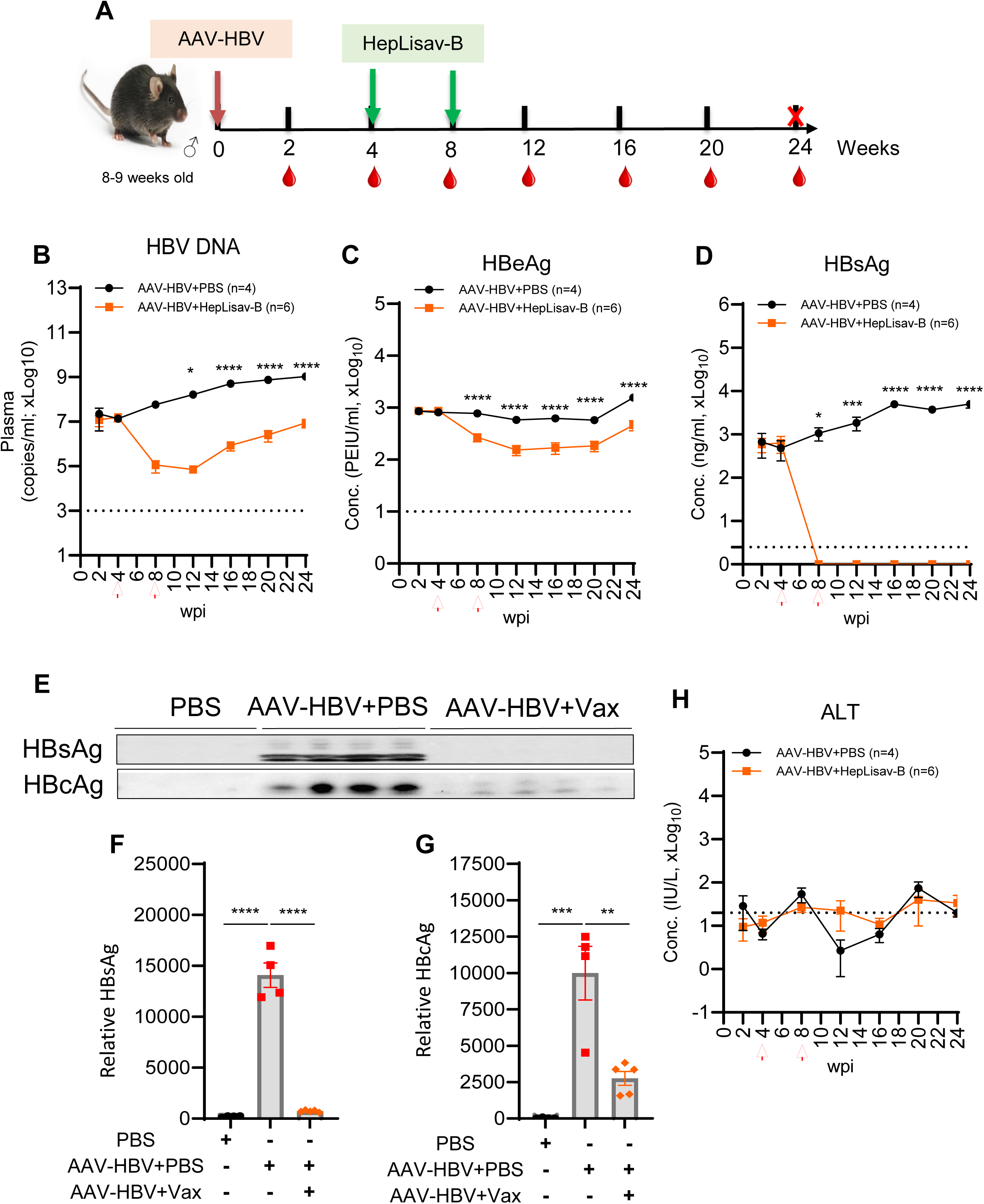
HEPLISAV-B suppresses HBV replication in chronic HBV mice. (**A**) Experimental design: C57BL/6 mice were inoculated with AAV-HBV and treated with PBS or HEPLISAV-B at 4- and 8-weeks post-infection (wpi). Blood was collected from 2–24 wpi; mice were euthanized at 24 wpi. (**B**) Plasma HBV DNA quantified by qPCR. Plasma (**C**) HBeAg and (**D**) HBsAg measured by ELISA. (E) Immunoblot analysis of plasma HBsAg and HBcAg at 24 wpi with (**F,G**) densitometric quantification. (H) Plasma ALT levels measured by ELISA. Data are mean ± SEM. Statistics: two-way ANOVA with Tukey post hoc test; *P<0.05, **P<0.005, ***P<0.0005, ****P<0.00005.

At termination, HEPLISAV-B markedly reduced hepatic HBV total RNA and 3.5 kb RNA (∼4-fold; Figure 2A–B), total intrahepatic HBV DNA (∼10-fold; Figure 2C), and AAV-HBV vector ITR DNA (∼4-fold; Figure 2D). Concordant reductions in hepatic HBs and HBc protein were confirmed by immunoblot and immunofluorescence (Figure 2E-J). Normalization to AAV-HBV template DNA showed proportional loss of HBV RNA and rcDNA (Figure S3A–C), consistent with non-cytolytic depletion of AAV-HBV episome DNA, a functional surrogate for HBV cccDNA in this model (Figure S3D).

**Figure 2.**
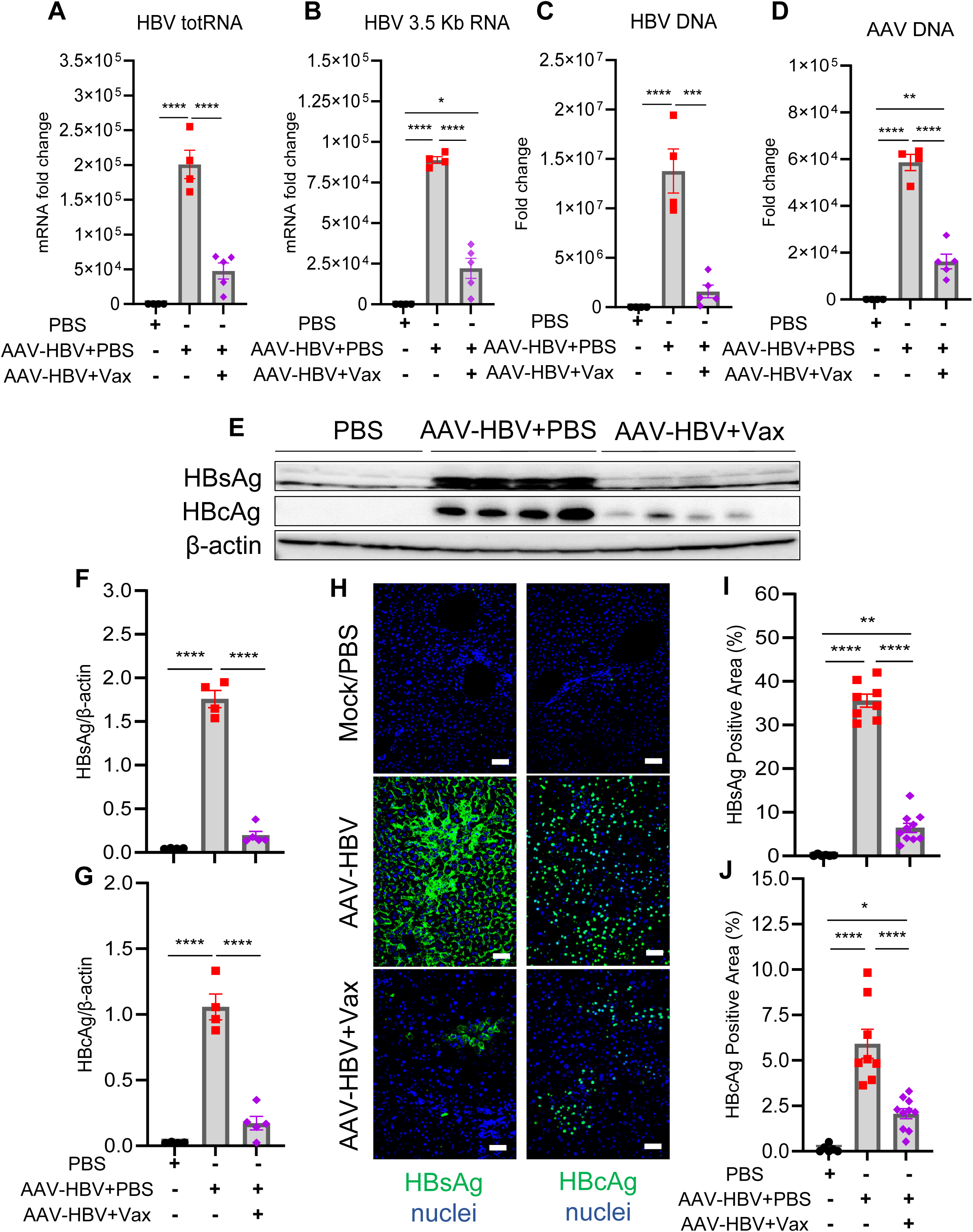
HEPLISAV-B reduces intrahepatic HBV RNA, DNA, and protein. (**A–D**) Intrahepatic levels of HBV total RNA, pgRNA (3.5 kb RNA), HBV DNA, and AAV inverted terminal repeat (ITR) DNA measured by RT-qPCR or qPCR at 24 wpi. (**E**) Immunoblot analysis of hepatic HBsAg and HBcAg with (**F,G**) quantification. (H) Immunofluorescence staining of intrahepatic HBsAg and HBcAg with (**I,J**) ImageJ quantification. Images acquired at 20x magnification. Data are mean ± SEM. Statistics: two-way ANOVA with Tukey post hoc test.

### HEPLISAV-B induces strong anti-HBs IgG1 and IgA responses in chronic HBV carrier mice

HEPLISAV-B elicited high-titer (≥10^4^ U/mL), durable anti-HBs IgG in chronically infected mice, comparable to naïve animals (Figure 3A). Vaccinated HBV-carrier mice displayed preferential induction of S-specific IgG1 and IgA (Figure 3B-D), indicating a Th2-skewed and mucosal-associated humoral profile.

**Figure 3.**
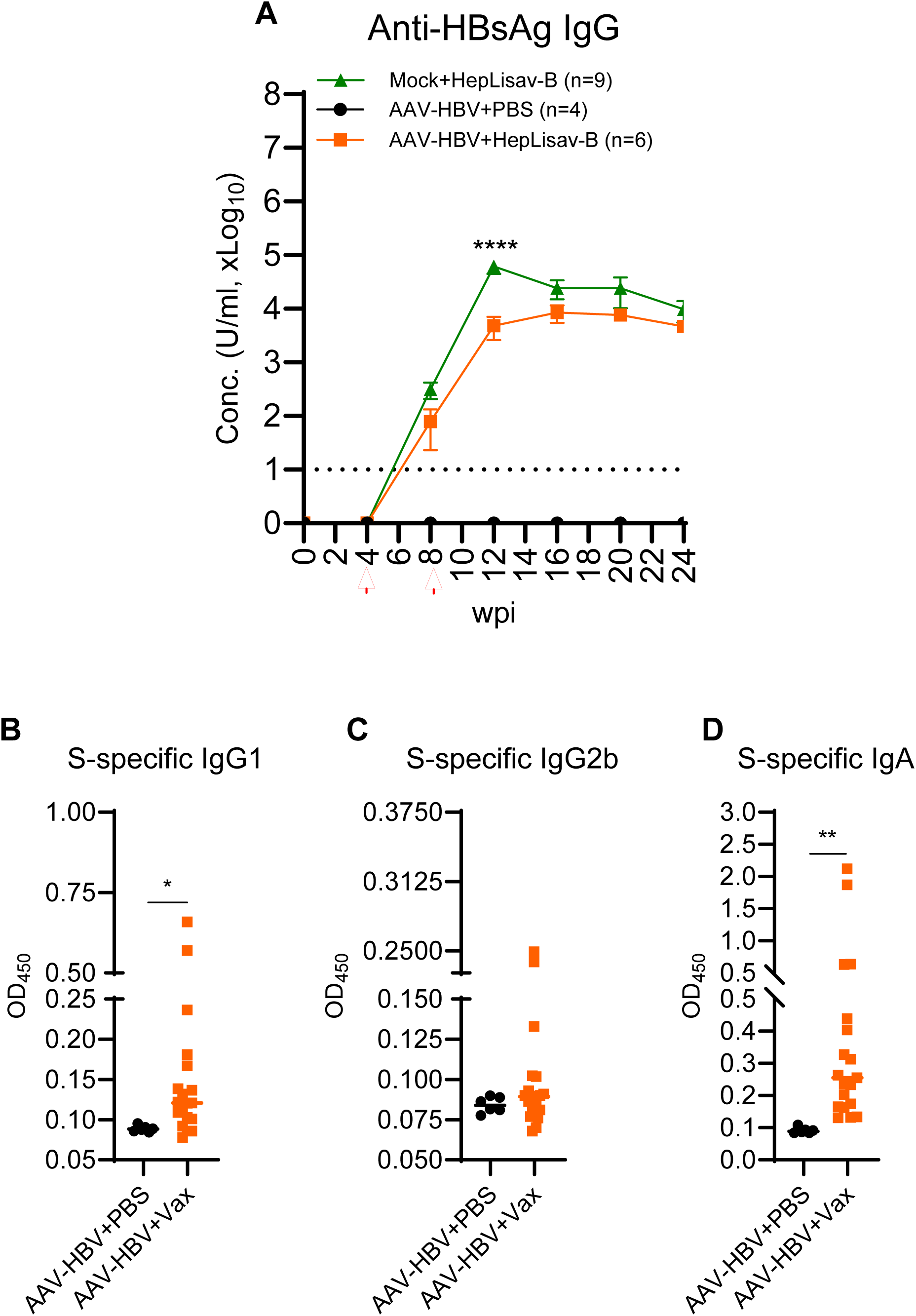
HEPLISAV-B induces robust anti-HBs humoral responses. (**A**) Plasma anti-HBs IgG levels measured longitudinally by ELISA (0–24 wpi). Plasma HBsAg-specific (**B**) IgG1, (**C**) IgG2b, and (**D**) IgA levels measured by ELISA. Data are shown as OD_450_. Bars represent median or SEM. Statistics: unpaired t test or two-way ANOVA with Tukey post hoc test.

Splenocytes from vaccinated HBV-carrier mice showed enhanced IFN-γ and TNF-α responses in CD4^+^ and CD8^+^ T cells upon stimulation (Figure 4). HBs-specific IFN-γ ELISPOT counts were increased (Figure 5A-B). Although total IgG ELISPOT numbers were unchanged (Figure 5C-D), HBs-specific IgG spot frequency and size were significantly greater, indicating higher-quality B-cell responses (Figure 5E-G). Together, these data show that HEPLISAV-B breaks HBsAg tolerance and restores antigen-specific T- and B-cell immunity.

**Figure 4.**
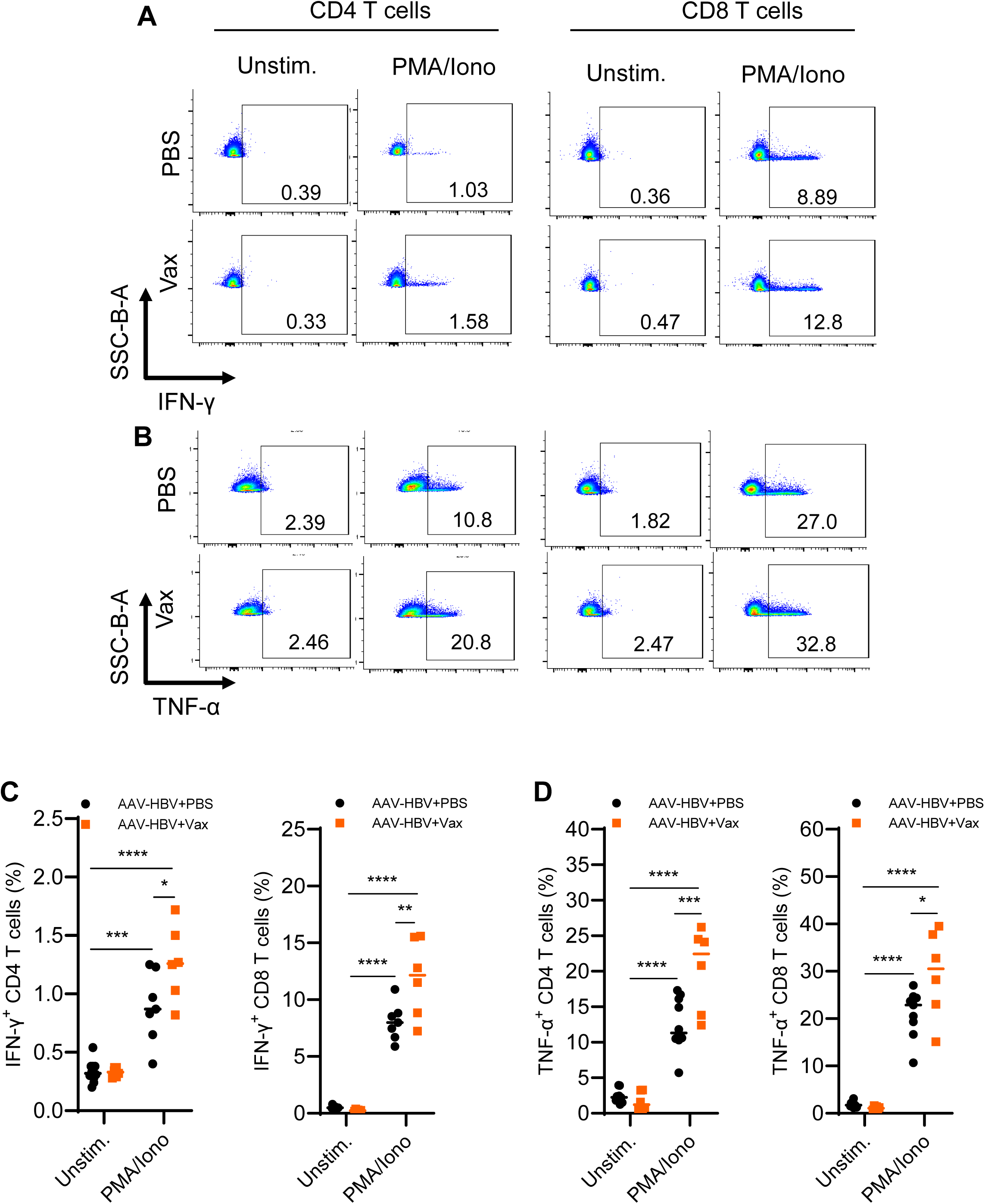
HEPLISAV-B enhances cytokine production by T cells. Splenocytes from chronic HBV mice treated with or without HEPLISAV-B were stimulated ex vivo with PMA/ionomycin in the presence of brefeldin A and analyzed by flow cytometry at 24 wpi. (**A**) Representative flow plots of IFN-γ⁺ and TNF-α⁺ CD4 and CD8 T cells. (**B–D**) Quantification of cytokine-producing T cells. Bars indicate median values. Statistics: two-way ANOVA with Tukey post hoc test.

**Figure 5.**
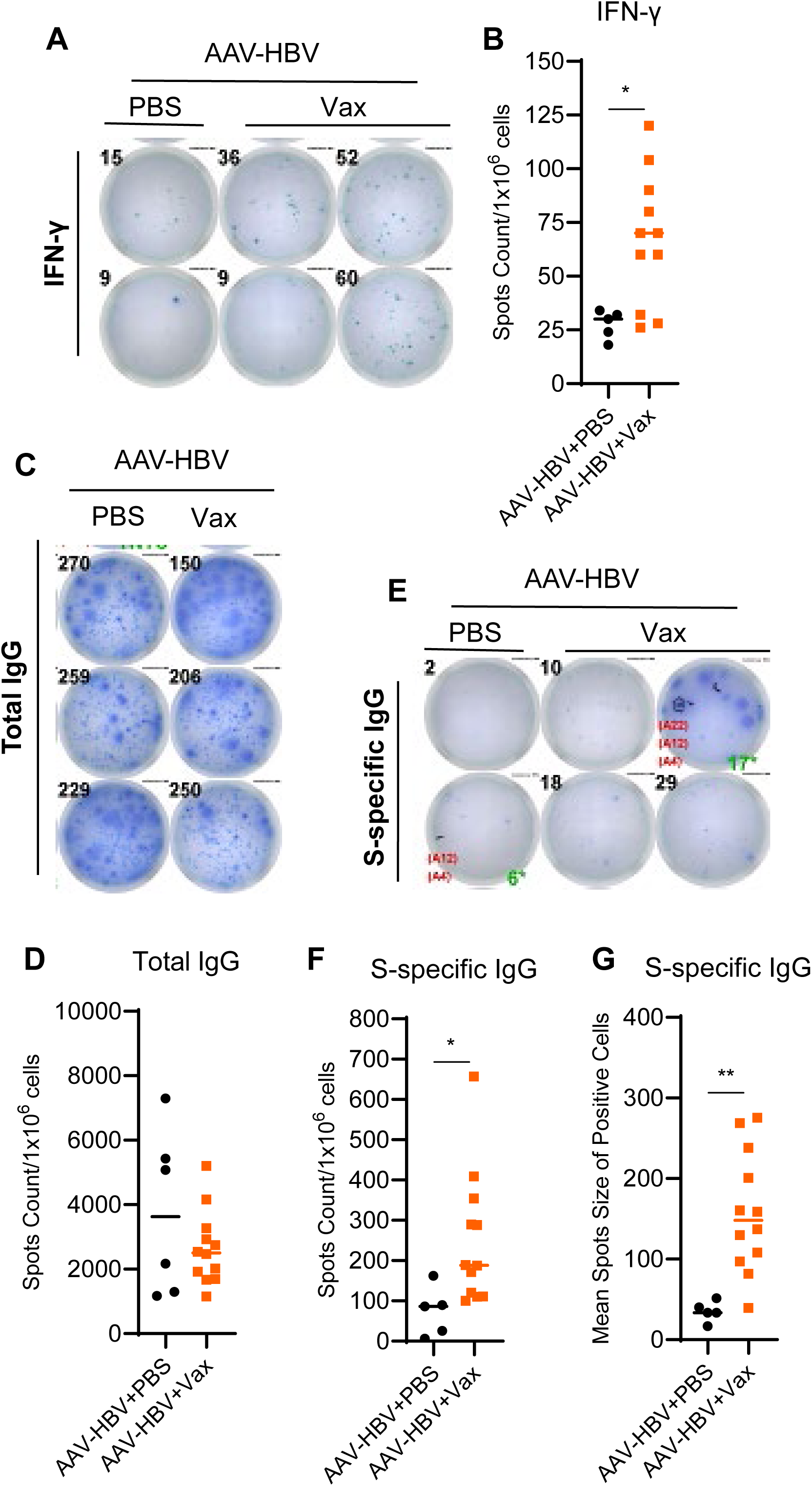
HEPLISAV-B restores HBsAg-specific T and B cell responses. (**A**) Representative IFN-γ ELISPOT wells following HBsAg stimulation with (**B**) summary quantification. (**C,D**) Total IgG ELISPOT wells and quantification. (**E**) Representative HBsAg-specific IgG ELISPOT wells with (**F,G**) spot counts and mean spot size. Data normalized per million live cells. Bars represent median values. Statistics: unpaired t test.

### Therapeutic efficacy of HEPLISAV-B requires CD4 T cells but not CD8 T cells

At 12 wpi, HEPLISAV-B increased liver and spleen size (Figure S4A-C) with enhanced intrahepatic immune infiltration but no increase in ALT or apoptosis (Figure S4D-E). Hepatic cytokine profiling revealed reduced IL-1β, IL-6, TGF-β, IL-21, and LY6D, with increased IL-27 and CXCL11 (Figure S5), suggesting diminished immunosuppression and enhanced T-cell recruitment.

Splenic analysis showed reduced frequencies of Tregs (CD25^+^FOXP3^+^ and CD25^-^FOXP3^+^; Figure S6A–B) and decreased PD-1 expression on CD4 T cells (Figure S6C-D), while CD8 PD-1 remained unchanged (Figure S6E). Memory profiling demonstrated increased TCM CD4^+^ and TEM CD8^+^ subsets (Figure S7A-F), indicating formation of functional memory compartments.

To define the cellular requirements for therapeutic activity, we depleted CD4 or CD8 T cells during vaccination (Figure S8A-D). CD4 depletion completely abolished virological control, preventing HBV DNA decline and HBsAg clearance (Figure 6A-B), and eliminating anti-HBs induction (Figure 6C). Correspondingly, reductions in hepatic HBV RNA, total DNA, and AAV-ITR DNA were lost (Figure 6D-F), and intrahepatic HBsAg/HBcAg remained high (Figure 6G-I). In contrast, CD8 depletion did not impair vaccine efficacy, and unexpectedly further increased anti-HBs IgG titers (Figure 6A-C), with preserved intrahepatic viral antigen clearance (Figure 6G-I).

**Figure 6.**
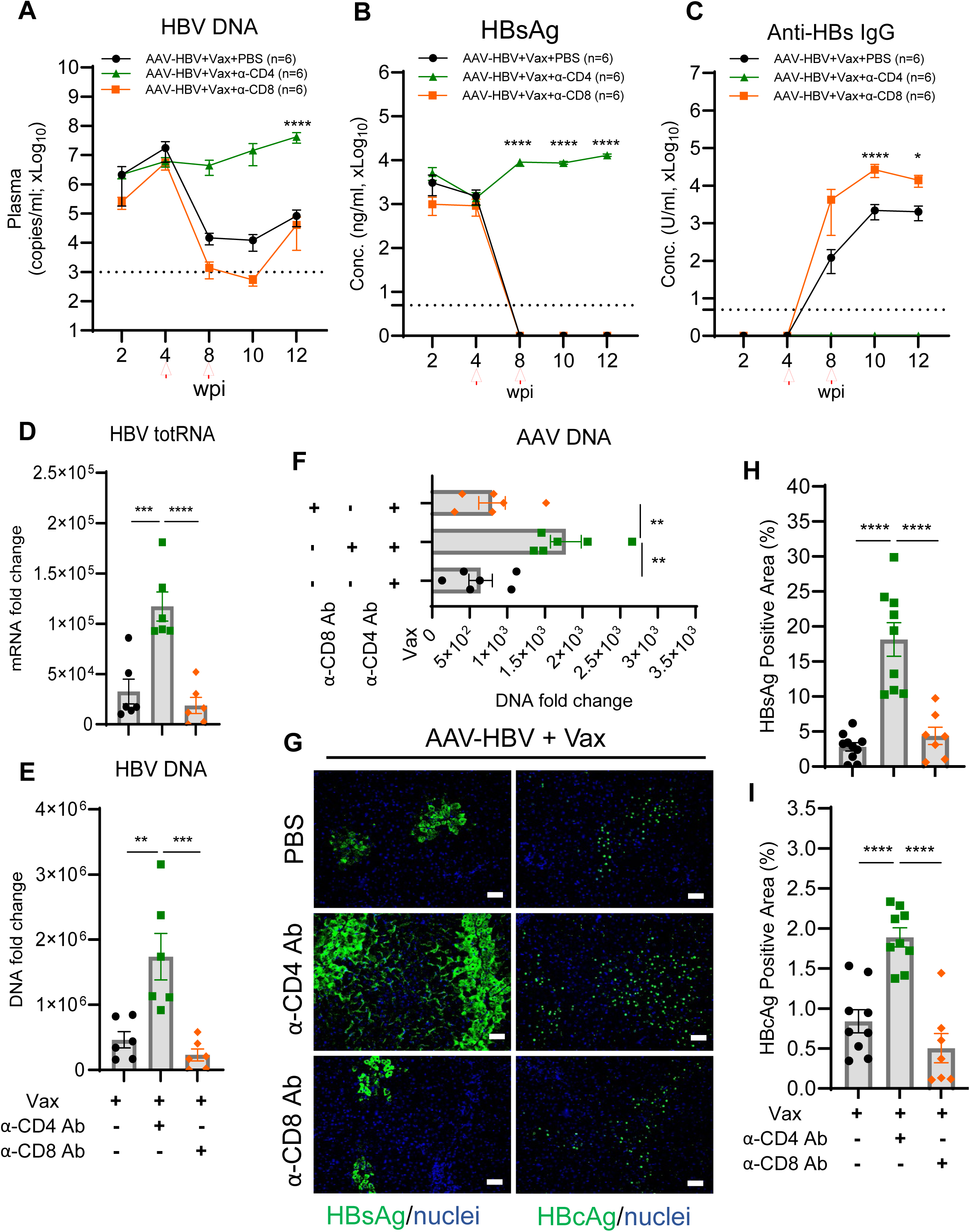
CD4 T cells are required for HEPLISAV-B–mediated HBV clearance. Chronic HBV mice treated with HEPLISAV-B received PBS, anti-CD4, or anti-CD8 antibodies. (**A**) Plasma HBV DNA by qPCR. Plasma (**B**) HBsAg and (**C**) anti-HBs IgG by ELISA. Intrahepatic (**D**) HBV total RNA, (**E**) HBV DNA, and (**F**) AAV DNA. (**G**) Immunofluorescence staining of hepatic HBsAg and HBcAg with (**H,I**) quantification. Data are mean ± SEM. Statistics: two-way ANOVA with Tukey post hoc test.

### Therapeutic efficacy of HEPLISAV-B is dependent on CD40/CD40L signaling

The interaction between CD40 on B cells and its ligand CD40L (CD154) on activated CD4 T cells is a central pathway mediating T cell help, driving B cell activation, clonal expansion, germinal center formation, and antibody production^25, 26^. To define the contribution of CD40/CD40L signaling to HEPLISAV-B–induced HBV suppression and antibody induction, HBV carrier mice were treated with an anti-CD40L blocking antibody in conjunction with HEPLISAV-B (Figure 7). Blockade of CD40/CD40L signaling completely abrogated HEPLISAV-B–induced HBsAg clearance, anti-HBs induction (Figure 7A-B), and hepatic reduction of HBs and HBc (Figure 7C-E), establishing the critical role of CD40/CD40L pathway in HEPLISAV-B–mediated antiviral immunity.

**Figure 7.**
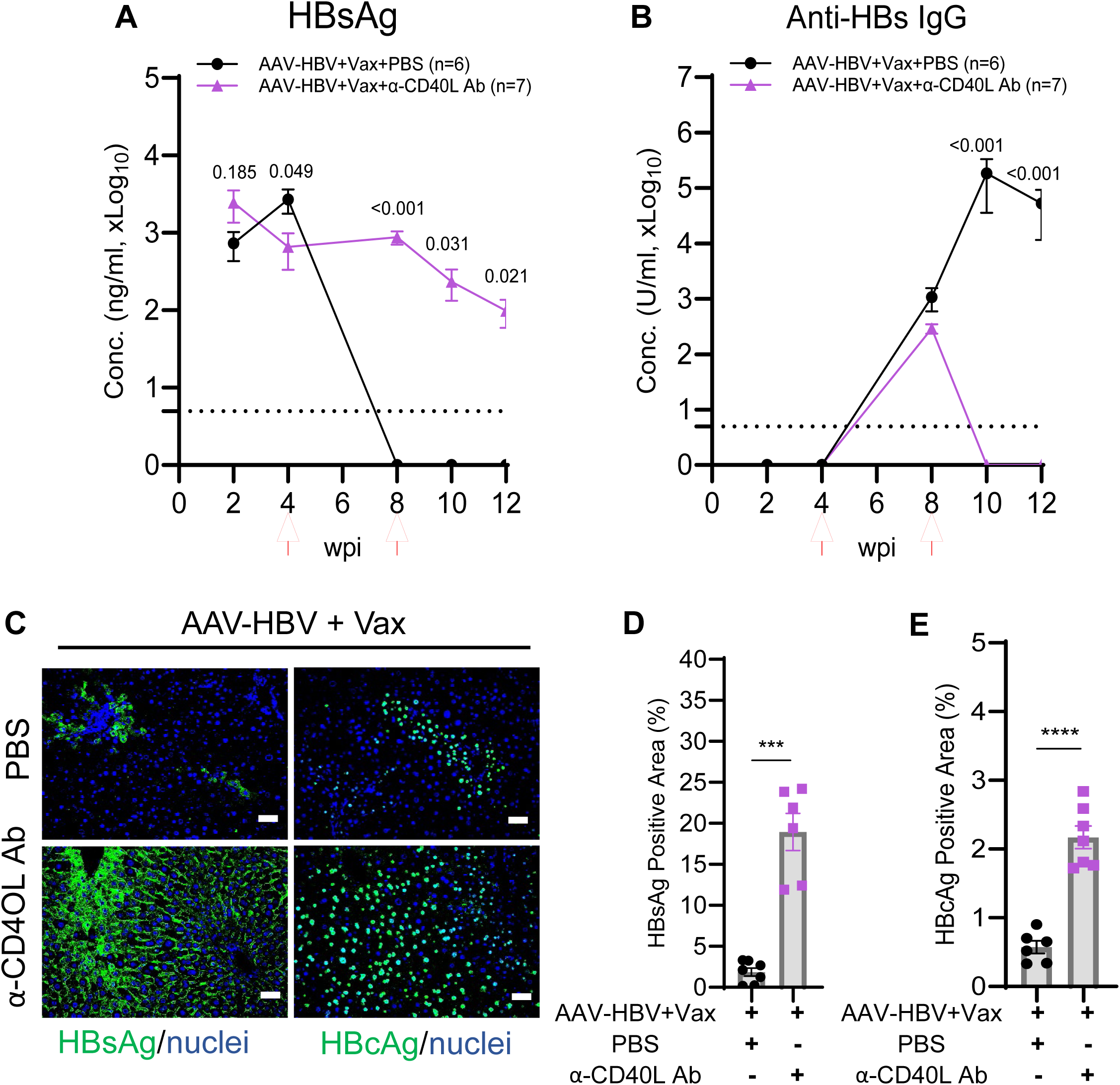
CD40/CD40L signaling is essential for HEPLISAV-B efficacy. HEPLISAV-B–treated chronic HBV mice received PBS or anti-CD40L antibody. Plasma (**A**) HBsAg and (**B**) anti-HBs IgG measured by ELISA. (**C–E**) Immunofluorescence staining and quantification of hepatic HBsAg and HBcAg. Data are mean ± SEM. Statistics: unpaired t test or two-way ANOVA with Tukey post hoc test.

## Discussion

We report that therapeutic vaccination with HEPLISAV-B (HBsAg/CpG-1018) overcomes HBV-induced immune tolerance and clears HBsAg in the AAV-HBV mouse model with high HBV viremia, outperforming alum- and Montanide/IL12-adjuvanted HBV vaccines. Virological control in the liver was non-cytolytic and accompanied by marked reductions in hepatic HBV transcripts and HBV DNA. Notably, AAV-HBV vector DNA (ITR-positive episomes; a cccDNA surrogate in this model) also declined by 10 folds, consistent with vaccine-driven clearance of HBV template DNA rather than hepatocyte injury. Interestingly, CD4 T cells, not CD8 T cells, were required for the HEPLISAV-B activity in HBV carrier mice. In addition, the CD40L/CD40 pathway is also critical.

HEPLISAV-B elicited a strong and durable anti-HBs response (≥10^4^ U/mL) with IgG1 and IgA enrichment, together with restoration of HBs-reactive T cells producing IFN-γ/TNF-α. At the systems level, the vaccine shifted the immune milieu from suppression/exhaustion towards effective help and memory: reduced Tregs and PD-1 on CD4 T cells, increased CD4⁺ central memory and CD8⁺ effector memory, and selective cytokine/chemokine profiles compatible with diminished regulatory^37^ and improved T-cell recruitment/function^38, 39^. These effects align with CpG-1018’s capacity to activate TLR9-expressing antigen-presenting cells and provide the necessary inflammatory context to overcome tolerance in chronic infection.

Mechanistically, the therapeutic efficacy required CD4 T cells and CD40/CD40L signaling. CD4 depletion abrogated HBsAg clearance, HBV DNA suppression, and anti-HBs induction, and prevented loss of intrahepatic HBsAg/HBcAg. In contrast, CD8 depletion enhanced anti-HBs titers, indicating CD8 T cells may suppress vaccine-mediated immune response and virological control via unknown mechanisms. Together, these data place the CD4 T/CD40L–CD40/B-cell axis at the core of HEPLISAV-B’s therapeutic effect and are consistent with recent observations that CD4 priming is pivotal for therapeutic vaccine efficacy. The transient rebound of circulating HBV DNA despite sustained HBsAg loss suggests persistence of non-productive nucleic acids (for example, defective particles or free DNA) and warrants further mechanistic work to define residual species and sources.

This study has several limitations. The AAV-HBV transduced mouse model lacks authentic HBV cccDNA with AAV-HBV episomal DNA as template to produce HBV in mouse hepatocytes. Thus, AAV episomes serve only as a surrogate for HBV template DNA loss. The exact identity and kinetics of residual circulating HBV DNA after vaccination remain unresolved, and the durability of control beyond the study window and the safety of repeated innate stimulation merit investigation. In mice, broad TLR9 expression in multiple immune cells is different from its restricted expression in B cells and pDCs. It will be of interest to define the specific cell types that directly respond to HEPLISAV-B in HBV carrier mice.

Despite these caveats, the data provide a clear mechanistic framework: CpG-1018-adjuvanted HBsAg restores effective CD4 help, licenses B-cell and T-cell immunity, and drives non-cytolytic clearance of viral products and template DNA, even with high-level viremia. To our knowledge, this is the first demonstration that HEPLISAV--B immunotherapy can deplete intrahepatic AAV-HBV template DNA via a CD4-dependent, non-cytolytic mechanism. These findings support clinical evaluation of HEPLISAV-B as a therapeutic vaccine in patients with high chronic HBV HBsAg/viremia.

## Acknowledgments

This work was supported by the National Institutes of Health (NIH) grants AI138797 and AI175800 (to L.S.), and by the National Cancer Institute - Cancer Center Support Grant (CCSG) – P30CA134274. We thank Weirong Yuan, Yichen Lu, and Sahra Sharifi—for their technical supports, and acknowledge the University of Maryland School of Medicine Animal Facility Core and the Histology and Flow Cytometry Shared Services of the University of Maryland Marlene and Stewart Greenebaum Comprehensive Cancer.

## Author contributions

Conceptualization: JA and LS. Methodology: JA and LS. Investigation: JA, JW and MF. Visualization: JA. JW and LS. Resources: LT, SK, LS. Supervision: LS. Writing—original draft: JA. Writing—review and editing: JA and LS.

## Supplemental Figures Legend

**Figure S1.**
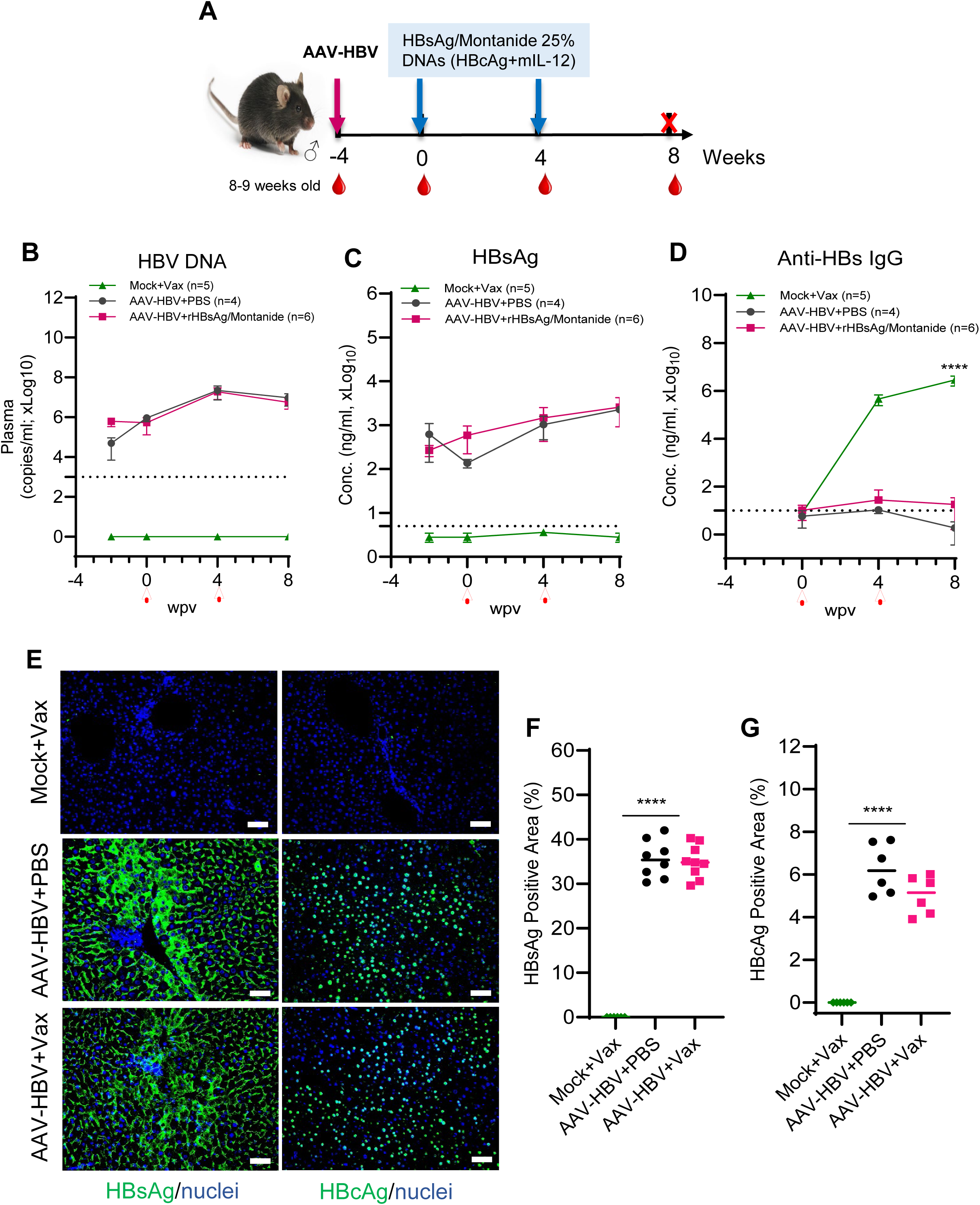
Conventional vaccination fails to break HBV immune tolerance. (**A**) Experimental design of Montanide-adjuvanted HBsAg plus DNA vaccination. (**B**) Plasma HBV DNA by qPCR. Plasma (**C**) HBsAg and (**D**) anti-HBs IgG by ELISA. (**E**) Hepatic immunofluorescence staining of HBsAg and HBcAg with (**F,G**) quantification. Images were acquired at 20x. Data are mean ± SEM. Statistics: two-way ANOVA with Tukey post hoc test.

**Figure S2.**
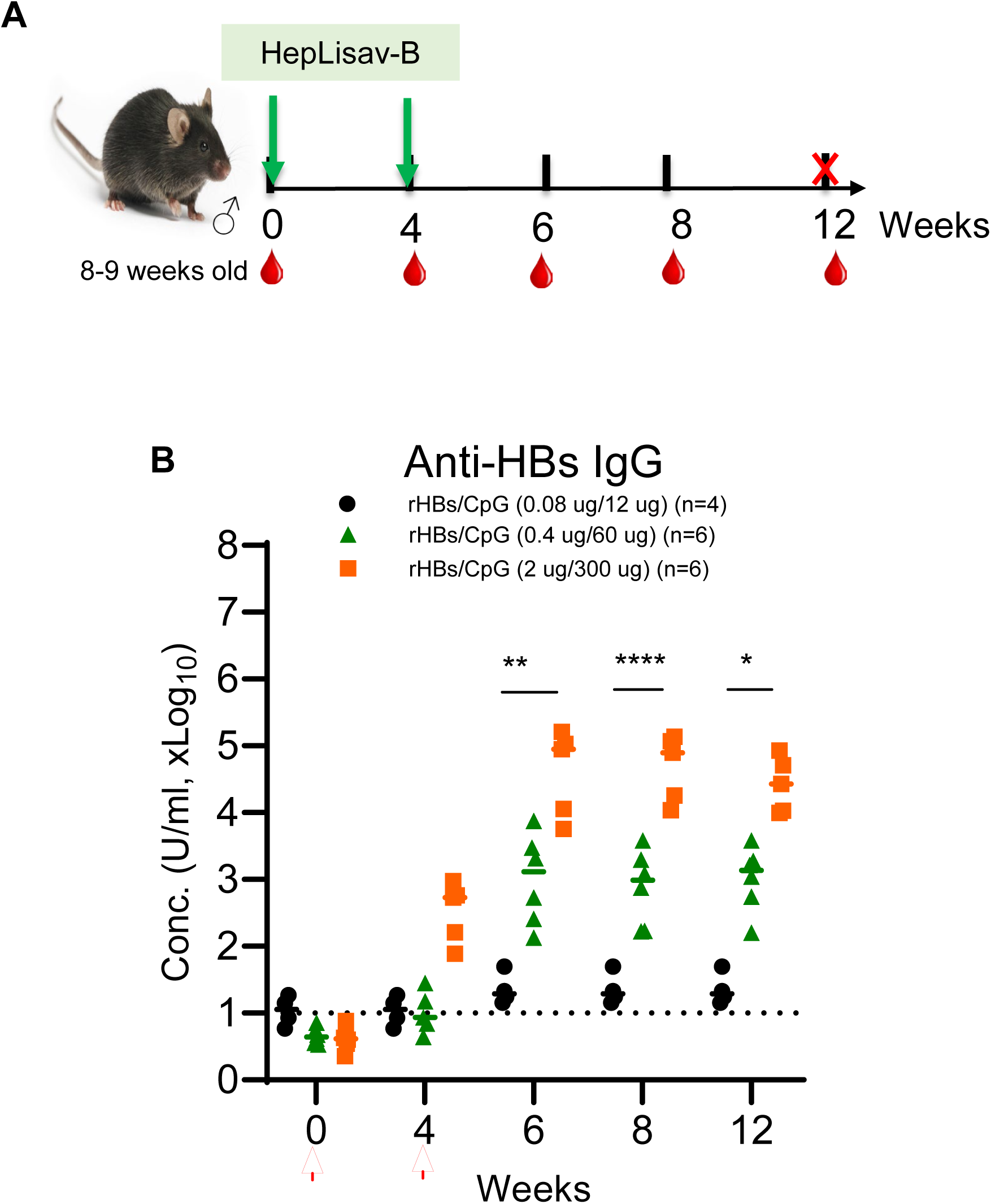
HEPLISAV-B dose titration in HBV-naïve mice. (**A**) Experimental design with three HEPLISAV-B dose levels. (**B**) Plasma anti-HBs IgG measured by ELISA over 12 weeks. Data are mean ± SEM. Statistics: two-way ANOVA with Tukey post hoc test.

**Figure S3.**
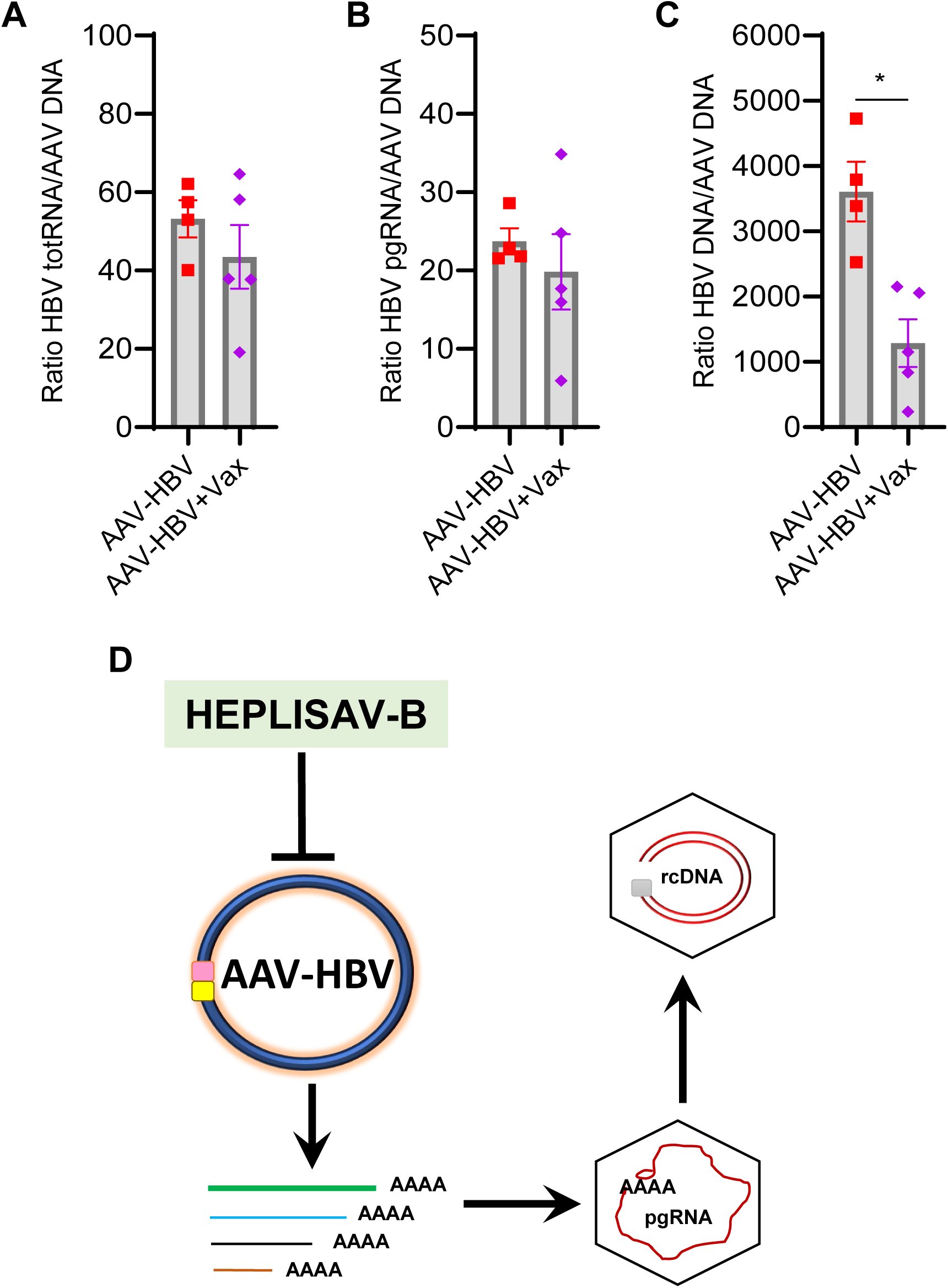
HEPLISAV-B reduces intrahepatic HBV templates. (**A–C**) Ratios of HBV RNA or DNA to AAV DNA in liver tissue. (**D**) Schematic illustrating HEPLISAV-B–mediated inhibition of HBV template activity and rcDNA generation. Bars indicate median values. Statistics: Mann–Whitney U test.

**Figure S4.**
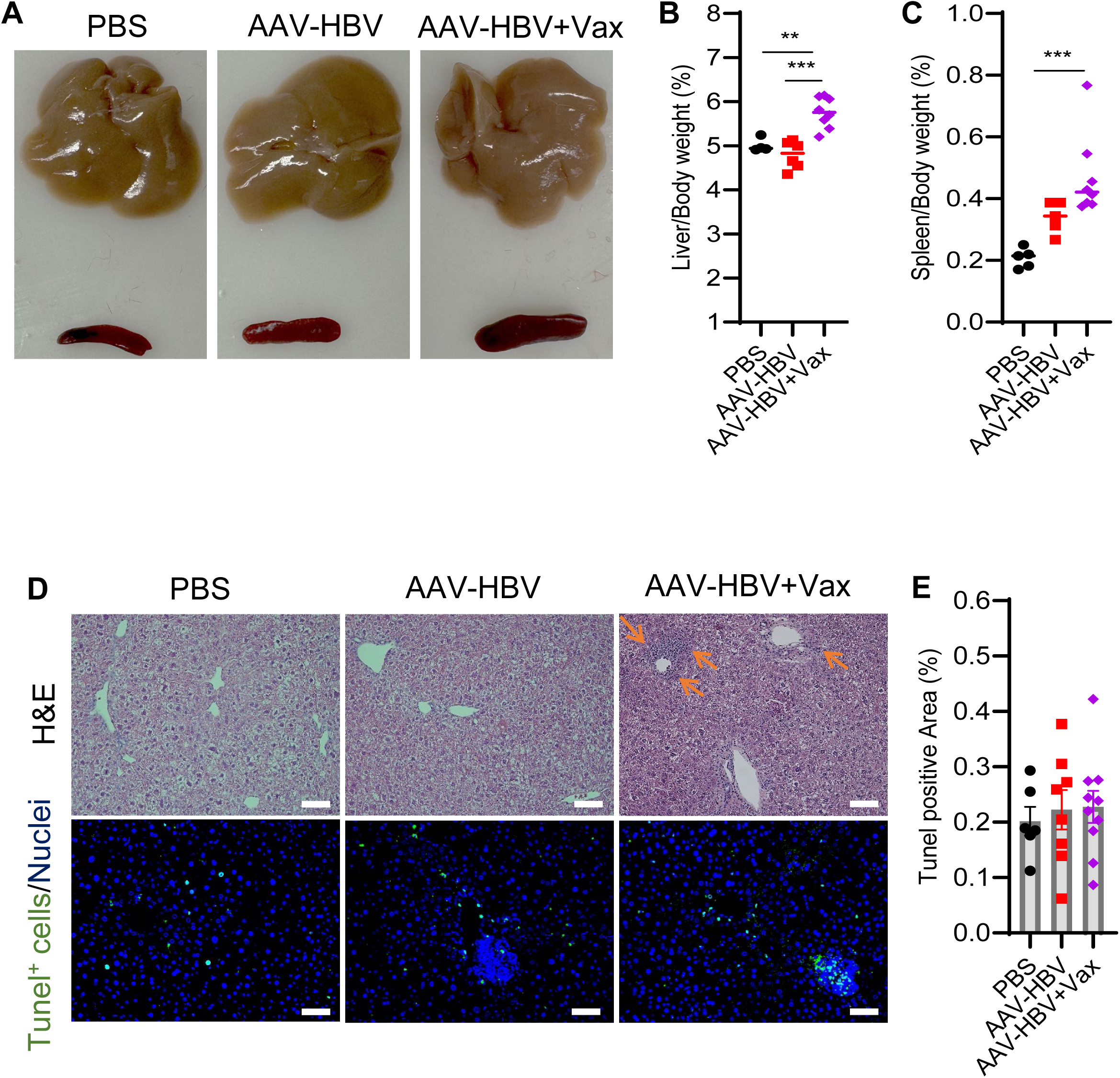
HEPLISAV-B induces non-cytolytic HBV clearance. (**A**) Representative liver and spleen images. (**B,C**) Organ-to-body weight ratios. (**D**) H&E and TUNEL staining of liver tissue with (**E**) quantification of TUNEL-positive area. Bars indicate median or SEM. Statistics: one-way ANOVA with Tukey post hoc test.

**Figure S5.**
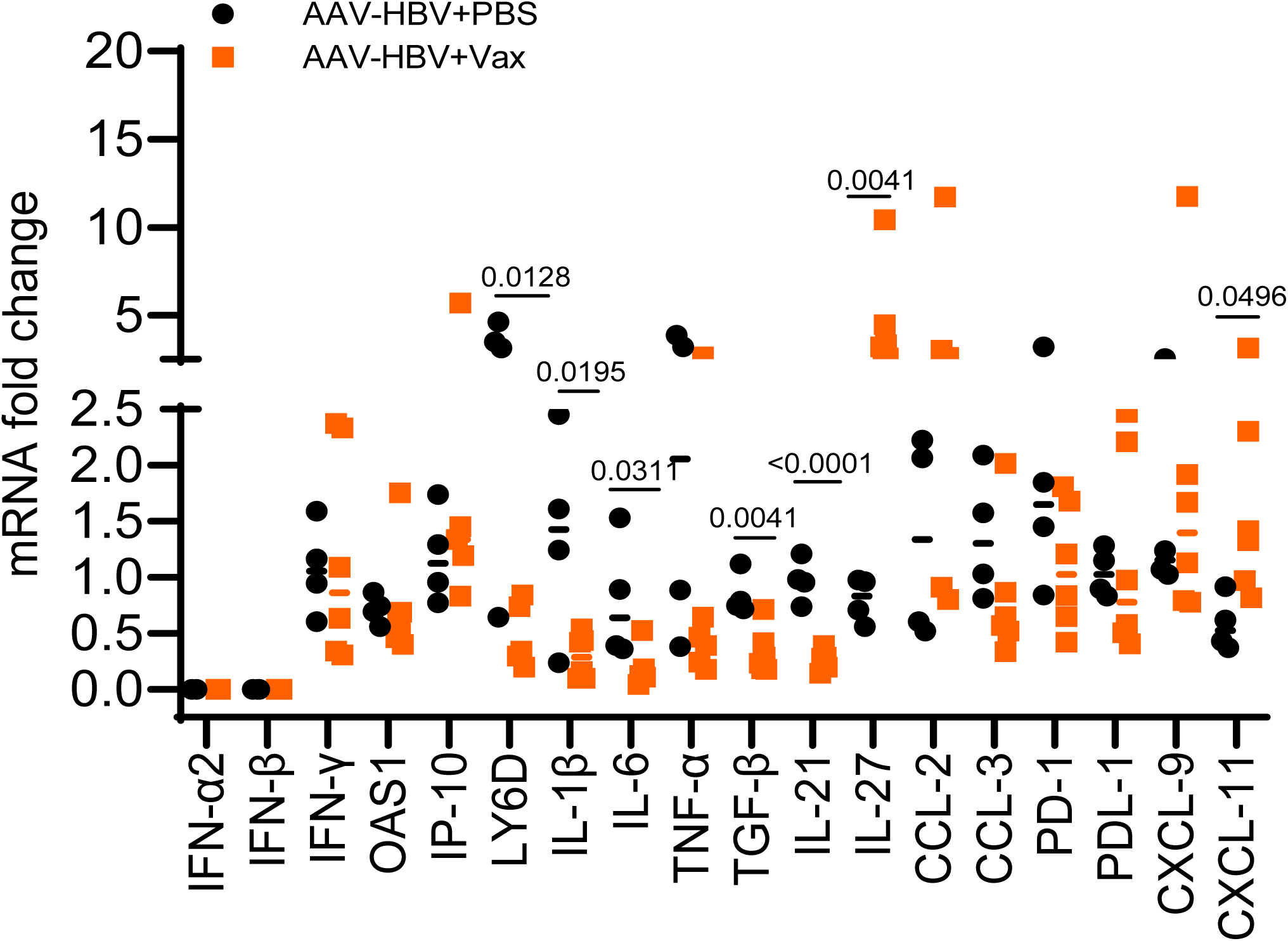
Altered intrahepatic cytokine and ISG expression following HEPLISAV-B. RT-qPCR analysis of hepatic interferons, ISGs, cytokines, chemokines, and fibrosis markers. Bars indicate median values. Statistics: unpaired t test.

**Figure S6.**
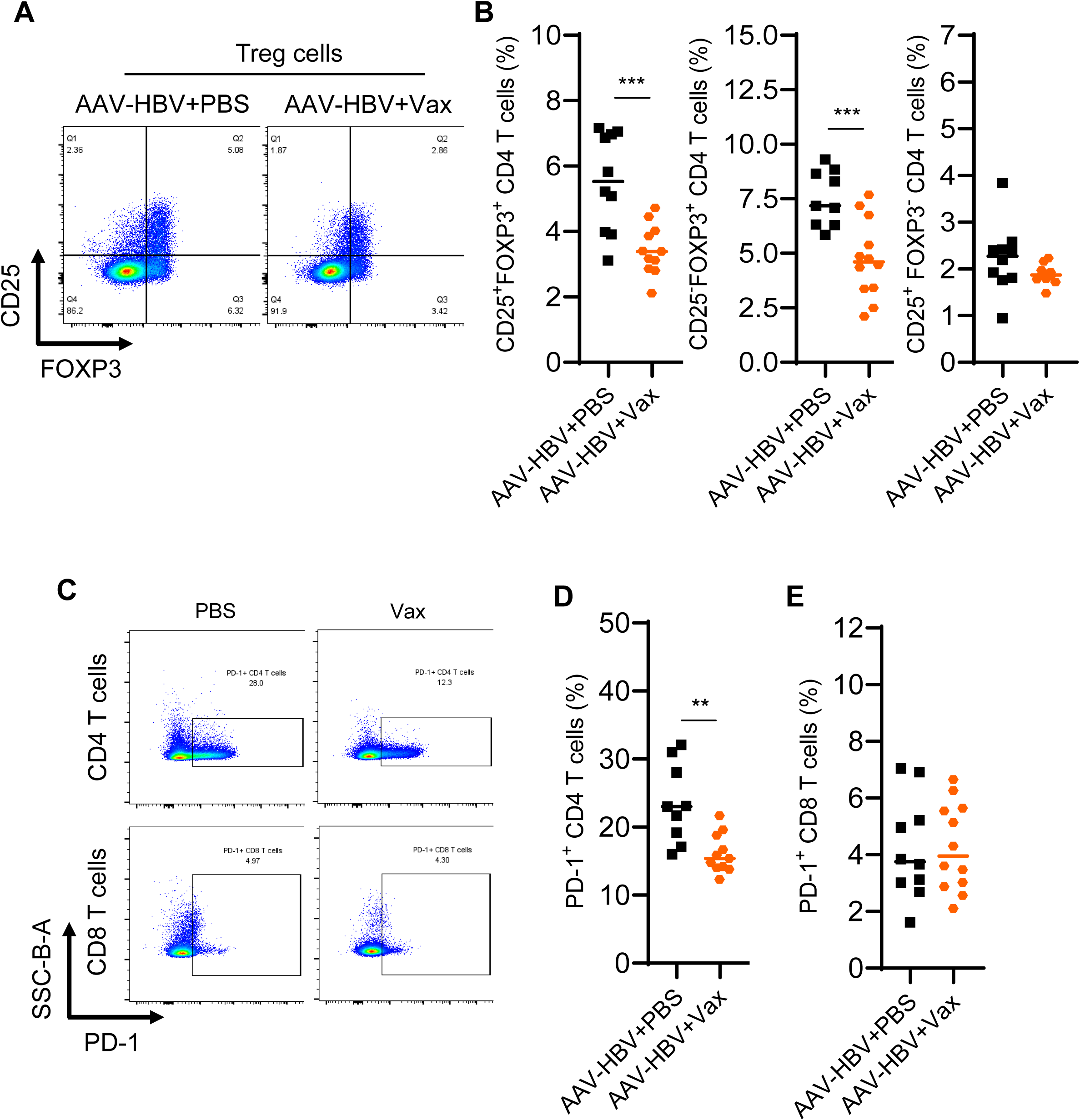
HEPLISAV-B reduces Tregs and PD-1 expression. (**A**) Representative flow plots of splenic CD25⁺FOXP3⁺ CD4 T cells. (**B**) Quantification of Treg subsets. (**C**) Representative PD-1 staining on CD4 and CD8 T cells with (**D,E**) summary data. Bars indicate median values. Statistics: unpaired t test.

**Figure S7.**
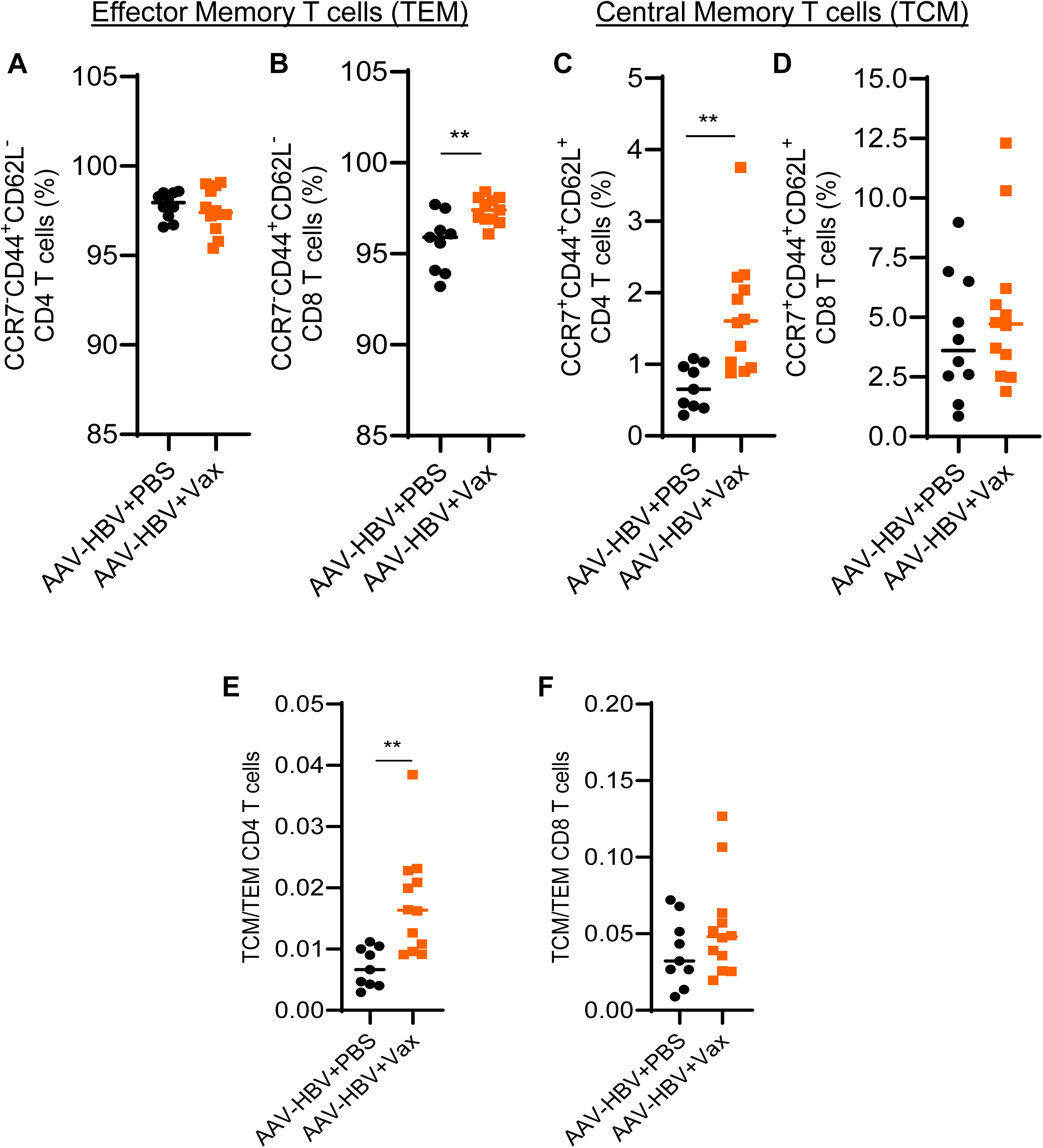
HEPLISAV-B promotes memory T cell differentiation. Quantification of splenic (**A,B**) effector memory (TEM), (**C,D**) central memory (TCM), and (**E,F**) TCM:TEM ratios for CD4 and CD8 T cells at 12 wpi. Bars indicate median values. Statistics: unpaired t test.

**Figure S8.**
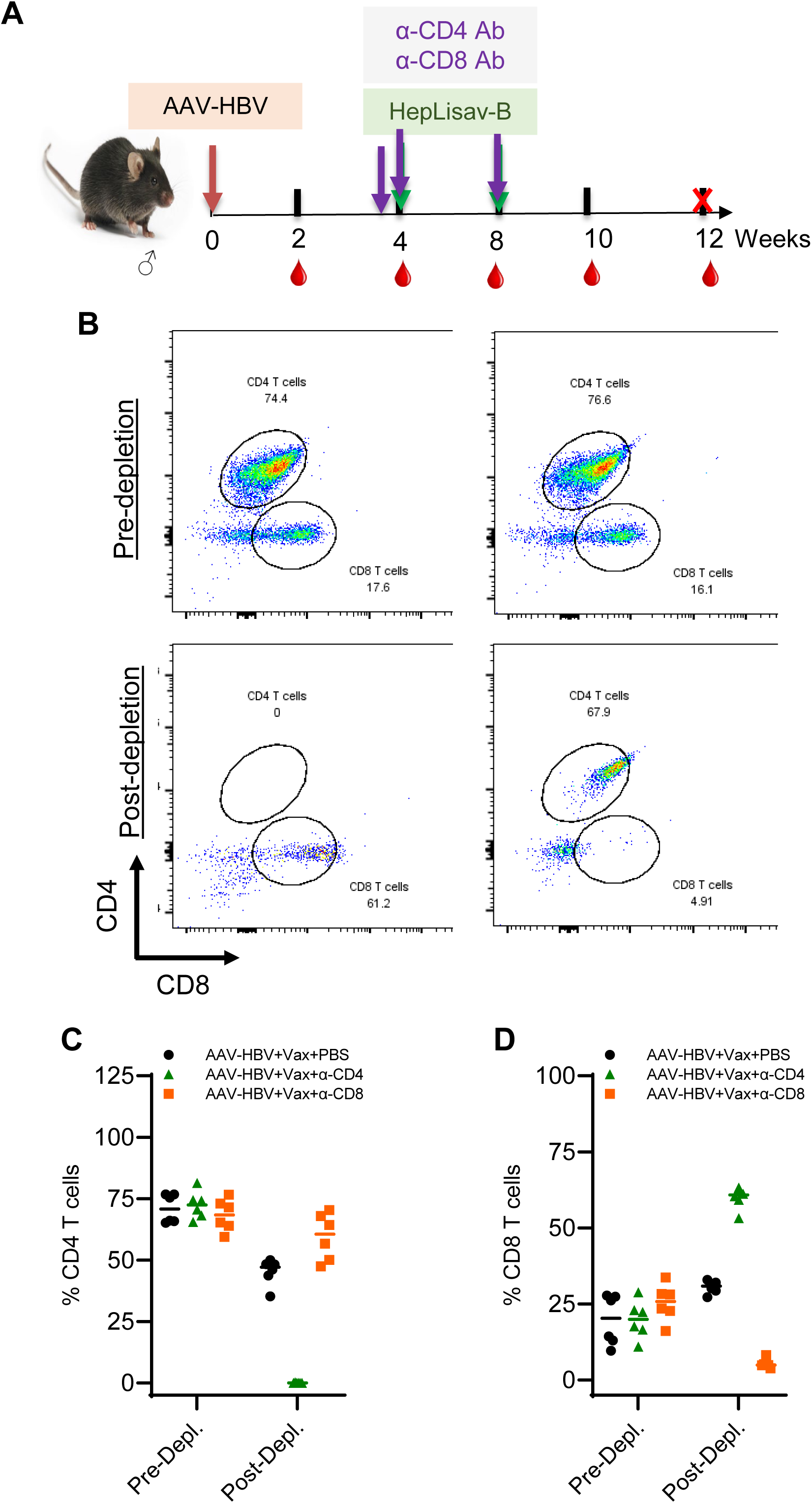
| Efficiency of CD4 and CD8 T cell depletion. (**A**) Experimental design of T cell depletion during HEPLISAV-B treatment. (**B**) Representative flow plots of CD4 and CD8 T cells. (**C,D**) Quantification of CD4 and CD8 T cell frequencies before and after depletion.

